# Seroprevalence of Hepatitis B and C among Healthcare Workers in Dutse Metropolis Jigawa State, Nigeria

**DOI:** 10.1101/327940

**Authors:** N.M. Sani, I. Bitrus, A.M. Sarki, N.S. Mujahid

**Affiliations:** Department of Microbiology and Biotechnology, Federal University Dutse Jigawa State-Nigeria; Department of Microbiology and Parasitology Aminu Kano Teaching Hospital Kano-Nigeria; Department of Medical Microbiology and Parasitology Bayero University Kano-Nigeria

**Keywords:** prevalence, hepatitis, viruses, healthcare workers, infection

## Abstract

Hepatitis is one of the neglected infectious diseases in sub Saharan Africa and most of the available data is based on blood donors. Health care workers (HCWs) often get infected as a result of their close contact with patients. A cross sectional study was conducted to determine the prevalence of hepatitis B and C among this group of professionals with a view to improving the quality of care to their patients. Hepatitis B and C infections pose a major public health problem worldwide. While infection is highest in the developing world particularly Asia and sub-Saharan Africa, healthcare workers are at higher risk of acquiring blood-borne viral infections, particularly Hepatitis B and C which are mostly asymptomatic. This study was aimed at determining the prevalence of Hepatitis B and C infections and associated risk factors among health care workers in Dutse Metropolis, Jigawa State - Nigeria. A standard rapid immuno-chromatographic technique i.e. rapid ELISA was used to screen all sera for Hepatitis B surface antigen (HBsAg) and Hepatitis C viral antibody (HCVAb) respectively. Strips containing coated antibodies and antigens to HBV and HCV respectively were removed from the foil. Strips were labeled according to samples. Using a separate disposable pipette, 2 drops of the sample (plasma) were added into each test strip and allowed to run across the absorbent pad. Results were read after 15 minutes. The prevalence of HBV and HCV infection in 100 healthcare workers was determined by testing the plasma collected from the clients during their normal checkup using HBsAg and HCVAb test strips. Results were subjected to statistical analysis using chi-square test. The prevalence of HBV among HCWs was 19 out of 100 (19.0%) and that of HCV was 5 out of 100 (5.0%) where in both cases, higher prevalence was observed among female nurses. It was also observed that all HCV positive cases were recorded among nurses only. The study revealed that nurses are at greater risk of contracting HBV and HCV due to their frequent contact with patients. It is therefore recommended that effective vaccination and other infection control measures be encouraged among healthcare workers.

## I. INTRODUCTION

**H**EALTH care workers in developing countries including Nigeria are at greater risk of infection from blood-borne pathogens particularly Hepatitis B (HBV) and Hepatitis C (HCV) viruses, due high prevalence of such pathogens in these countries [1].

There are about 130 million persons having hepatitis C infection worldwide. Its prevalence varies from region to region. The highest prevalence (15–20%) has been found in Egypt while the United Kingdom has lowest prevalence (0.01 –0.1%), [2]. According to an estimate, 27% of cirrhotic patients and 25% of patients with hepatocellular carcinoma are infected by HCV [3].Hepatitis B on the other hand, has been found to infect about 350 million people globally. Like hepatitis C, the prevalence of hepatitis B also varies from low to high in different parts of the world. Both HBV and HCV are blood-borne hepatotropic viruses but have distinct routes of transmission. Most commonly, HBV is acquired by vertical transmission from an infected mother or via horizontal transmission in childhood [4].

However, HCV is primarily transmitted parenterally in adulthood by intravenous drug use, blood transfusion, or medically related parenteral exposures, but rarely through the placenta, breastfeeding, or sexual contact. Previous studies by [5] have shown that frequent parenteral injections with non-disposable needles were highly associated with HCV seropositivity and that most of the anti HCV positive persons did not have a history of blood transfusion or intravenous drug use, the most commonly documented risk factors for HCV. Hepatitis C virus is less efficiently transmitted by single small dose percutaneous exposures [6]. Organ transplant from infected donors is also a known cause of viral transmission. In Pakistan several previously conducted studies have shown different prevalence rates of HBV and HCV infection.13–19 [2] Approximately three million health care workers (HCW) are exposed to percutaneous blood – borne virus each year. It is estimated that 66,000 hepatitis B virus (HBV) and 16,000 hepatitis C virus (HCV) are acquired annually [7] infections are important risk factors for hepatocellular carcinoma and other liver related morbidity [8]. The HBV carrier rate varies widely from 0.01% to 20% in different geographical regions of the world [9]. The prevalence of HBV in some Middle East countries ranges between 2% to 7% in the general population, while HCV is much lower in these countries [10]

The HCW including clinicians, nurses, laboratory technicians, other hospital technicians, administrators and cleaning staff are exposed to an increased risk of occupational infection with hepatitis B and C virus [11]. In tropical countries the risk of occupational transmission of these viruses is further increased by the excessive handling of needles and lancets to test for common tropical diseases. For example, more than 100 million tests for malaria are performed each year [11] Data exists concerning the epidemiology of HBV and HCV in several study populations from Sudan [8]. However we are unaware of published data on these viruses among HCW in the country. It is known that hepatitis B (HBV) and C viruses (HCV) are among the most frequent blood borne pathogens. Occupational exposure to them refers mainly to healthcare workers because of frequent contact with body fluids[12]. The risk of transmitting an infection following an occupational exposure depends on lots of factors and is on average 0.5% for HCV and 18–30% for HBV [13]. In United States it’s been estimated that a prevalence of HBV infection among healthcare workers (HCWs) was approximately 10 times higher than among the general population [14]. Although the risk of being infected with HBV is greater than HCV, there are efficient means of pre- and post-exposure prophylaxis (hepatitis B immunoglobulin and/or hepatitis B vaccine – obligatory for health care workers). According to the World Health Organization, 5% of HCWs in central part of Europe are exposed to at least one sharp injury contaminated with HBV per year and 1.7% – to the one contaminated with HCV [15]. Polish Chief Sanitary Inspectorate estimated that there were about 1513 new cases of hepatitis B in 2009 in Poland and there are 450 000 people infected with HBV. Implementing obligatory vaccinations against HBV in children and adolescents in1994 immunized individuals up to the age of 18 against this virus. In the vaccinated group acute hepatitis B is rarely observed [16]. Hepatitis C morbidity is dissimilar to the one caused by HBV; the extent of years is increasing, although in 2007–2009 this trend was revised. It’s estimated that there are about 730 000 people infected with HCV with a morbidity rate of 7–8 cases per 100 000 inhabitants [17]. The most often genotypes recognized in Poland are: 1b (75%), 3a (17%) and 1a (about 5%) [18]. All employees in Poland are obliged to undergo pre-employment and periodic medical examinations. The only laboratory tests that are compulsory for workers being at risk of exposure to HBV and HCV (according to the Polish guidelines on prophylactic examinations) to acquire are total serum bilirubin and alanine aminotransferase (ALT). Serologic tests aimed at early detection of HBV and HCV infection are facultative. As an employer covers the costs of prophylactic care, performing these tests quite often depends on the amount of money the healthcare setting wants to spend on laboratory tests. In Poland, there were a total of 2933 occupational diseases diagnosed in 2010. Recently, there are around 30–45 occupational cases of hepatitis B and 99–119 of hepatitis C recognized per year, mostly in healthcare workers [19].

Hepatitis B virus, abbreviated HBV, is a species of the genus Orthohepadnavirus, which is likewise a part of the hepadnaviridae family of viruses [20]. This virus causes the disease hepatitis B [21]. In addition to causing hepatitis, infection with HBV can lead to cirrhosis and hepatocellular carcinoma [1]. It has also been suggested that it may increase the risk of pancreatic cancer [21]. Hepatitis C virus (HCV) is a small (55–65 nm in size), enveloped, positive-sense single-stranded RNA virus of the family Flaviviridae. Hepatitis C virus is the cause of hepatitis C and some cancers such as Liver Cancer (Hepatocellular carcinoma abbreviated HCC) and lymphomas in humans [22]. The hepatitis C virus belongs to the genus Hepacivirus, a member of the family Flaviviridae. Until recently it was considered to be the only member of this genus. However a member of this genus has been discovered in dogs canine Hepacivirus [23]. There is also at least one virus in this genus that infects horses [24]. Several additional viruses in the genus have been described in bats and rodents. [23]. Hepatitis C virus has a positive sense single-stranded RNA genome. The genome consists of a single open reading frame that is 9,600 nucleotide bases long [25]. This single open reading frame is translated to produce a single protein product, which is then further processed to produce smaller active proteins.

At the 5’ and 3’ ends of the RNA are the UTR that are not translated into proteins but are important to translation and replication of the viral RNA. The 5’ UTR has a ribosome binding site (IRES - Internal ribosome entry site) that starts the translation of a very long protein containing about 3,000 amino acids [26].

The core domain of the hepatitis C virus (HCV) has IRES containing a four-way helical junction that is integrated within a predicted pseudo knot. [27]. The conformation of this core domain constrains the open reading frame’s orientation for positioning on the 40S ribosomal subunit. The large pre-protein is later cut by cellular and viral proteases into the 10 smaller proteins that allow viral replication within the host cell, or assemble into the mature viral particles [24]. Structural proteins made by the hepatitis C virus include Core protein, E1 and E2; nonstructural proteins include NS2, NS3, NS4A, NS4B, NS5A, and NS5B.

The proteins of this virus are arranged along the genome in the following order: N terminal-core-envelope (E1)–E2–p7-nonstructural protein 2 (NS2)–NS3–NS4A–NS4B–NS5A– NS5B–C terminal. The mature nonstructural proteins (NS2 to NS5B) generation relies on the activity of viral proteinases [28]. The NS2/NS3 junction is cleaved by a metal dependent autocatalytic proteinases encoded within NS2 and the N-terminus of NS3. The remaining cleavages downstream from this site are catalyzed by a serine proteinases also contained within the N-terminal region of NS3.

The core protein has 191 amino acids and can be divided into three domains on the basis of hydrophobicity: domain 1 (residues 1–117) contains mainly basic residues with two short hydrophobic regions; domain 2 (resides 118–174) is less basic and more hydrophobic and its C-terminus is at the end of p21; domain 3 (residues 175–191) is highly hydrophobic and acts as a signal sequence for E1 envelope protein [28].

Both envelope proteins (E1 and E2) are highly glycosylated and important in cell entry. E1 serves as the fusogenic subunit and E2 acts as the receptor binding protein. E1 has 4–5 N-linked glycan and E2 has 11 N-glycosylation sites.

The p7 protein is dispensable for viral genome replication but plays a critical role in virus morphogenesis. This protein is a 63 amino acid membrane spanning protein which locates itself in the endoplasmic reticulum. Cleavage of p7 is mediated by the endoplasmic reticulum’s signal peptidases. Two transmembrane domains of p7 are connected by a cytoplasmic loop and are oriented towards the endoplasmic reticulum’s lumen [28].

The risk of infection by (HBV) and (HCV) among health care personnel is a major public health issue, especially in low incomes countries. Indeed, healthcare providers are exposed to different types of infections where one of the main microbial reservoirs may be the patient. The infection can be directly transmitted by the patient or indirectly through contact with blood, body fluids or equipment [29]. The present study was conducted to determine the seroprevalence of HBV and HCV among healthcare workers with a view to assessing the risk factors for the acquisition of these infections in the study population.

### II MATERIALS AND METHODS

### A. Study Site

The study was conducted in three separate hospitals within Dutse metropolis namely; Rasheed Shekoni Specialist Hospital (RSSH), Dutse General Hospital (DGH) and Police Hospital Dutse (PHD), Dutse Local Government Area, Jigawa state-Nigeria.

### B. Study Participants

The participants were healthcare workers (Doctors, Nurses and Medical Lab. Scientist) from the three hospitals in Dutse Metropolis. The sample size was determined using the formula described by [30] and was found to be 100.

### C. Sampling Technique

A random sampling technique was used to collect data from health care workers that met the set inclusion/exclusion criteria and were therefore recruited for the study.

### D. Informed Consent

To each health care worker, a copy of consent form was administered and duly signed by the participant accordingly. Information on selection criteria, procedure and benefits of the study were explained to them. They were also made to understand that their participation was voluntary, and they were free to decline at any point.

### E. Questionnaire Administration

A structured closed ended questionnaire designed for the study was used to obtain data on demographic and social status of the participants as well as the risk factors for HBsAg and HCV infection. Parameters captured include: age, gender, Profession and marital status after which five milliliter (5ml) of whole blood was collected from each participant using sterile syringe and needle. The samples collected were centrifuged at 3,000 rev/min to get the plasma which was stored at refrigeration temperature before use.

### F. Assay Procedure

Samples were brought to room temperature by thawing and strips containing coated antigen and antibody to HBV HCV respectively were removed from the foil. Strips were labeled according to samples. Using a separate disposable pipette, 2 drops of each sample (plasma) was added into a test strip and allowed to run across the absorbent pad. Results were read after 15 minutes. Two controls were used in the test, one red band indicating positive, one band designating a negative test and no band at all for an invalid result. [31].

|  | Gender |  |  |
| --- | --- | --- | --- |
|  | F | M | Total |
| Negative | 31(38.3%) | 50(61.7%) | 81 |
| Positive | 10(24.4%) | 9(15%) | 19 |
| Total | 41(41.0%) | 59(59.0%) | 100% |

|  | AGE/YEARS |  |  |  |  |
| --- | --- | --- | --- | --- | --- |
|  | <25 | 25-30 | 31-40 | 41-50 | Total |
| Negative | 4(4.9%) | 34(42.0%) | 37(45.7%) | 6(7.4%) | 81 |
| Positive | 0(0.0%) | 7(17.1%) | 10(21.3%) | 2(25.0%) | 19 |
| Total | 4(4.0%) | 41(41.0%) | 47(47.0%) | 8(8.0%) | 100% |

### G. Principle of the Test

This is a qualitative lateral flow chromatographic immunoassay. When an adequate volume of the test specimen is dispensed onto the sample pad of the test device, the specimen migrates by capillary action across the strip. The antibody or antigen if present in the specimen binds with its corresponding conjugated antigen or antibody on the test device (strip) thereby forming a reddish coloration in the presence of a chromogen.

### H. Statistical Analysis

The data was subjected to Pearson’s Chi-Square test using Statistical Package for Social Sciences (SPSS) version 22.

## III RESULTS

Table 1 shows the distribution of Hepatitis B Virus infection different age groups of the subjects screened. The highest number of positive cases was found among the group aged 41-50 years with 2(25.0%) positivity, while the least was found among those <25 years.

**Table 1. Prevalence of HBV among different groups**

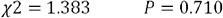

The distribution of HBV with respect to gender is shown in Table 2. It is clear from the table that out of 100 clients, 9(15%) males were positive while 41 females representing 41% were positive. From the table, the prevalence of hepatitis B virus was higher in female clients.

**TABLE II.**

|  | BLOOD GROUP. |  |  |  |  | Total |
| --- | --- | --- | --- | --- | --- | --- |
|  | 0- | 0+ | A+ | AB | B |  |
| Negative | 1(1.2%) | 52(64.2%) | 6(7.4%) | 11(13.6%) | 11(13.6%) | 81 |
| Positive | 0(0.0%) | 13(20.0%) | 4(40.0%) | 1(8.3%) | 1(8.3%) | 19 |
| Total | 1 | 65 | 10 | 12 | 12 | 100 |

**PREVALENCE OF HEPATITIS B VIRUS AMONG DIFFERENT GENDER GROUPS**

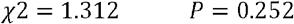

Table 3 shows the distribution of hepatitis B among blood groups. High prevalence of hepatitis B was recorded among A^+^ and O^+^ which was found to be 4(40.0%) and 13(20.0%) respectively with low prevalence in blood group O^-^.

**TABLE III PREVALENCE OF HEPATITIS B AMONG BLOOD GROUPS**

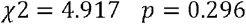

Figure 1 shows the distribution of HBV based on profession; the highest number of positive cases was recorded among Nurses with 11 (28.9%) while the least was recorded among Pharmacist and Doctors respectively. These categories of health care workers by nature of their exposure always had close contact with patients and body fluids.

**Fig. 1.**
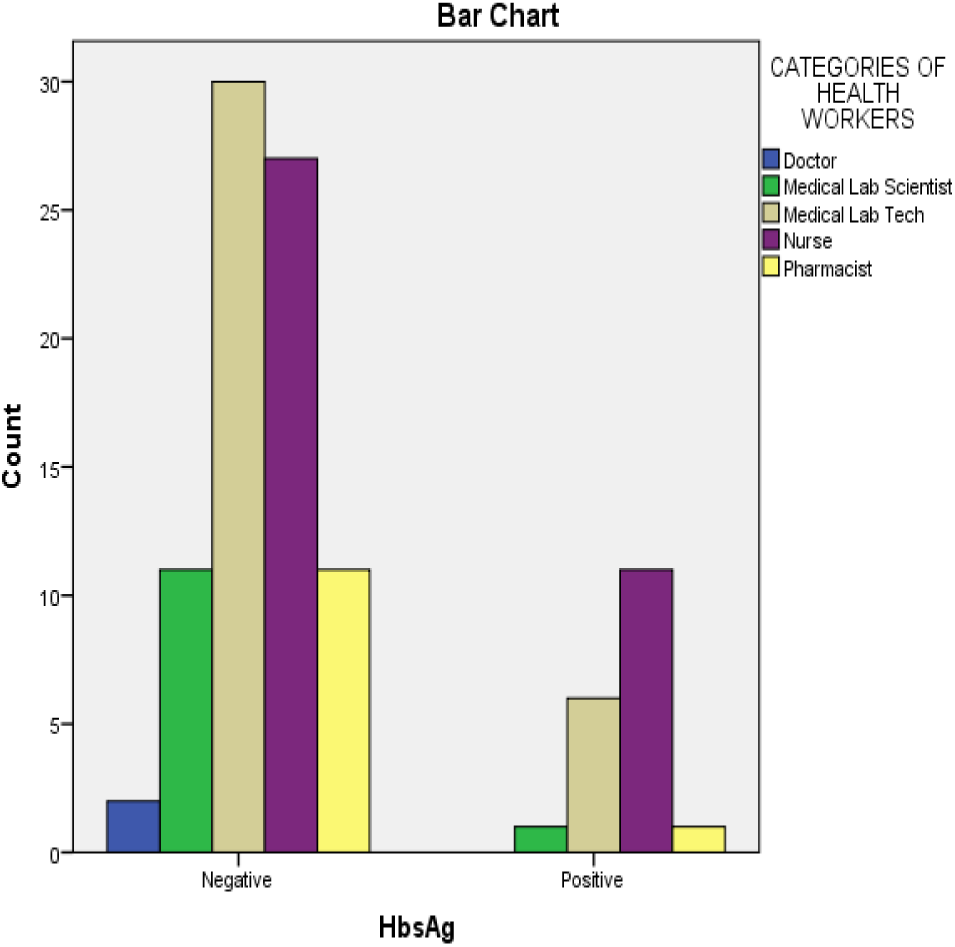
Prevalence of Hepatitis B among different categories of health care workers.

Table 4 shows the distribution of Hepatitis C Virus infection among different age groups of the subjects screened. The highest number of positive cases was found among the group aged 30-40 years with 2(40%) positivity, while the least was found among those <25 years.

**TABLE IV.**
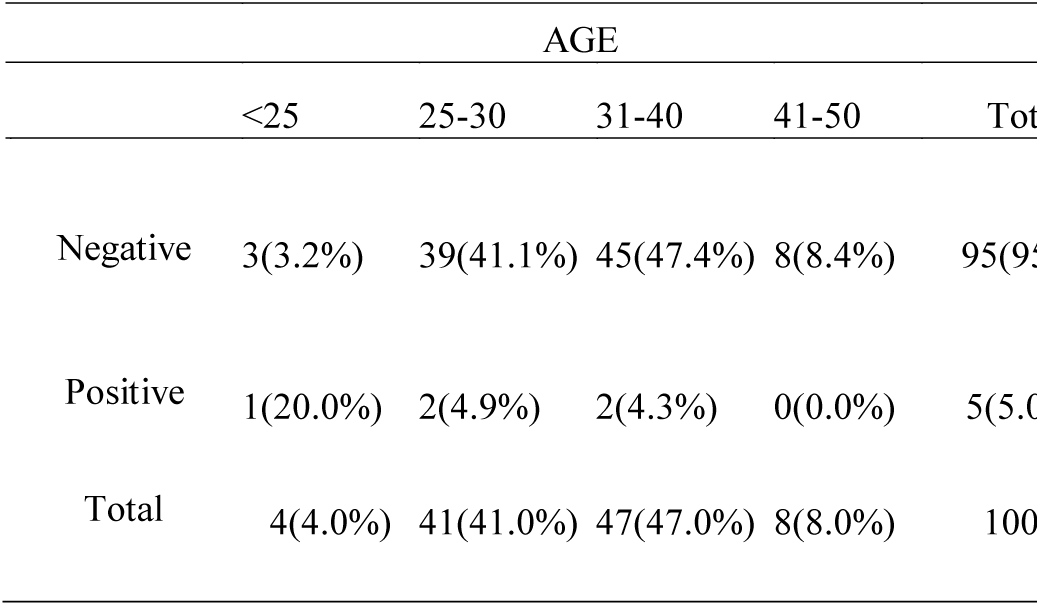
HEPATITIS C VIRUS (HCV) IN RELATION TO AGE.

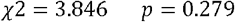

Figure 2 shows the prevalence of HCV based on the categories of HCWs. The highest number of positive cases was recorded among Medical Lab.Tech. 3(60 %) and Nurses with 2(40 %) while none were recorded among Pharmacist, medical lab scientist and Doctors respectively.

**Fig. 2.**
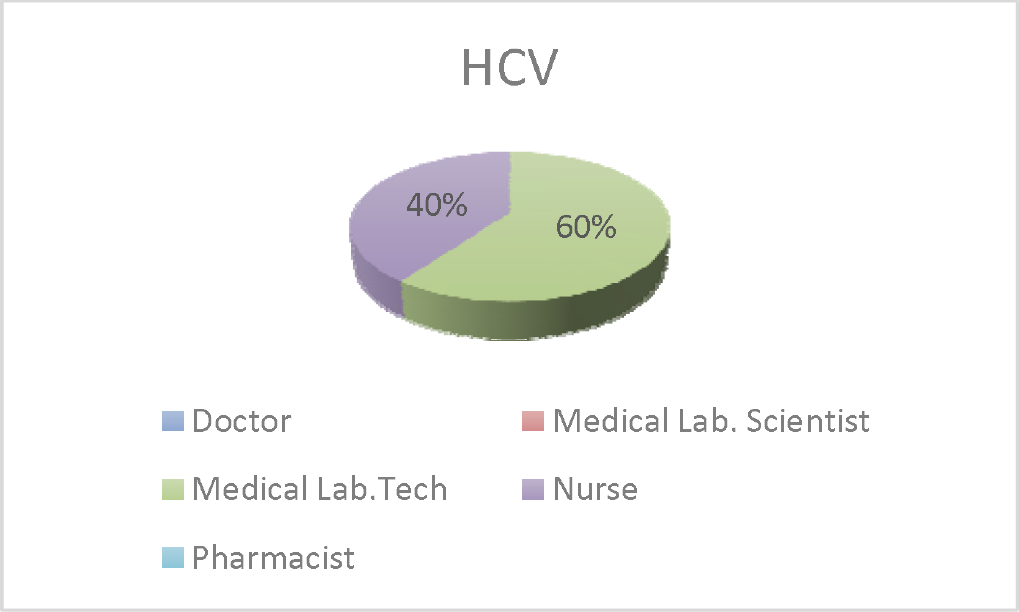
Prevalence of HCV based on Professional Category.

Table 5. Shows the prevalence of hepatitis C among blood groups. Higher prevalence of this virus was found among O^+^ 5(7.7%) with low prevalence in blood group O^-^, A^+^, AB and B. This can be as a result of the facts that, blood group O are universal donor while the others are not and as such 0% prevalence was recorded among the other groups.

**TABLE V.** PREVALENCE OF HEPATITIS C VIRUS (HCV) AMONG DIFFERENT BLOOD GROUPS OF HCWS.

|  | BLOOD GROUPING |  |  |  |  | Total |
| --- | --- | --- | --- | --- | --- | --- |
|  | O<sup>-</sup> | O<sup>+</sup> | A<sup>+</sup> | AB | B |  |
| Negative | 1(1.1 %) | 60(63.2%) | 10(10.5%) | 12(12.6%) | 12(12.6%) | 95(95.0 %) |
| Positive | 0(0.0 %) | 5(7.7%) | 0(0.0%) | 0(0.0 %) | 0(0.0 %) | 5(5.0%) |
| Total | 1(1.0 %) | 65(65.0%) | 10(10.0 %) | 12(12.0%) | 12(12.0%) | 100(100.0 %) |

## IV DISCUSSION

Occupational exposure of HBV is a well-recognized risk for health care workers (HCWs) [32] In developing regions, 40% - 65% of HBV infections in health-care workers occurred due to per-cutaneous occupational exposure [33]. Risk of HBV infection is primarily related to the degree of contact with blood in the work place and also to the hepatitis B-e antigen (HBeAg) status of the source person. The result obtained from this study showed a prevalence of 19.0% which indicates a high prevalence of Hepatitis B virus among heal h care workers at the locations of study. According to the findings of [34], the common factor for those at high risk has been high frequency of contact with patients’ blood [34]. In a study conducted among surgeons in Lagos Nigeria by [35], the prevalence of HBsAg among this category of Health workers was 25.7%. However in contrast, HBsAg seropositivity of 17.8% and antiHBs positivity of 79.2% were recorded among hospital workers in Senegal which agrees with the result obtained in this study. From a similar study conducted among health care workers in Uganda, a prevalence of 9.0% was recorded for current infection [36]. Prevalence of HbsAg recorded at the location of this study could also be that the prevalence of HBV infection is increasing, this is particularly so in view of the inadequate enlightenment on the infectious nature of the virus. Considering various categories of the HCWs, the nurses screened had a prevalence of 28.9% which was the highest prevalence among the health workers screened, compared to the Medical Laboratory Scientists with a prevalence of 8.3%. Laboratory technicians, recorded a prevalence of 16.9%. These category of health care workers by nature of their exposure always had close contact with patients and body fluids. This probably accounted for the high prevalence rates among these subjects. Similar observations have been reported in other studies with laboratory technicians, dentists and nurses being disproportionately affected [37] [38]. The variations among the different cadres of HCWs might be a reflection of the different levels of risk of exposure to a hazardous work environment the different categories of health care workers operate in. Based on gender, male subjects recorded a prevalence of 15% while females recorded 24.3%. The difference between these gender groups was not found to be statistically significant (p=0.252).Other studies however, showed the same results while others found that males are more infected than females [39], although there is no clear explanation of such variation among these studies, the nature of the society being modest may make gender of little or no influence on the spread of HBV infection. However gender seems not to be a relevant factor in this work.

With regards to the age of subjects screened, different studies concerning the prevalence of HBV infection among health care workers including this studies, reflects a trend of age related seropositivity, it was found that subjects aged 31-40 years recorded a higher prevalence, this is in agreement with the findings of [40] who observed that those who are aged over 30 were significantly higher than those who aged less than 30 years. This is in an agreement with other studies carried out by other investigators, [41] who showed hat the incidence of HBV markers increased with age and duration of employment. This however, may reflect the higher risk of exposure in the corresponding age in the general population. The prevalence among other health personnel showed that those with high proximity to blood and body fluids may be at greater risk, this is evident in the low percentage recorded among pharmacists who by nature of their work do not handle blood and body fluids as much as do scientists, Technicians and nurses. [42], made similar findings when HBV prevalence among Nurses, Medical Laboratory Scientists, Doctors, Dentists and Ward Orderlies were compared with other category of health workers. Differentials in knowledge about the dangers of hepatitis B and C virus infection and the available prevention strategies might also partly explain the observed differences in prevalence. It is worthy of note that individuals without any knowledge on the infectious nature of neither the virus nor its prevention had a higher risk of life time exposure to hepatitis B virus infection. The high prevalence of HBV among the health personnel probably reflects acquiring HBV infection during the performance of their duty schedules. [43] noted that since HBV can survive in dried blood specimen for a long time, the possibility that health workers can be infected even in situation less likely could not be ruled out.From this study, the overall seroprevalence of HCV among the health care workers was found to be 5.0 per cent. This figure is comparable to previous reports of [44] where the recorded 4% prevalence in India. The seroprevalence of HCV in the general population has been studied extensively and reports from different parts of India show the seroprevalence of HCV infection to be as varied as 0.3 to 11.3 per cent [45] [46] [47]). But most of the studies have shown the prevalence of HCV to be less than 2 per cent in the general population. In our study, the seroprevalence of HCV in health care workers was found to be considerably higher among Medical Lab.Tech. 3(8.3%) and Nurses 2(5.3%), as seen in table 5. The high prevalence of HCV among health care workers may be due to their exposure to infected blood/blood products of patients of HCV infection [44]. This increased exposure may be in the form of accidental needle-pricks, contact of cut skin surface with blood/blood products, improper disposal of infected medical waste, etc.

The results of this study support the prevailing evidence for HCV as an occupational hazard to health care workers. This view is supported by the report of [48], which indicated that the prevalence of HBV and HCV among health care workers is related to their work. The finding of a high prevalence of HBV infection among health care personnel is a major concern not only regarding the continuous spread of the infection, but the fact that health workers are daily exposed to the risk of infectious diseases in the course of their work. Considering other possible risk factors, it was observed that those who had blood transfusion recorded a higher prevalence compared to those who had none. This highlighted the risk involved in the transfusion of unscreened blood. The practice of giving unscreened blood to patients should therefore be avoided in view of the possibility of HBV infection. The World Health Organization [49] and [50] strongly recommend screening of blood meant for transfusion for infectious agents such as HBV, HCV and HIV. However, [51] commented that after controlling other variables, longer duration in service remained significantly associated with a lower risk of current infection. This finding is at variance with what has been reported in other studies, [36].

## CONCLUSION

A relatively high positivity rate was found among HCW. These results were in agreement with previous reports affirming that HCWs are at risk for infection with both HBV and HCV via exposure to infected blood or body fluids.

The prevalence of current hepatitis B virus infection and life time exposure to hepatitis B virus infection among health care workers was high. Exposure to potentially infectious body fluids was also high and yet only a small percentage of HCWs are vaccinated against hepatitis B virus infection. Considering the risk of transmitting HBV to patients, there is an urgent need to focus efforts on reducing transmission through improving the work place environment and ensuring prompt vaccination of all health care workers who are highly susceptible to the infectious virus. It is also important that health personnel be properly informed about their risk of HBV infection, so as to adopt measures to avoid been infected. This will help to maintain the integrity of the health system by protecting its workforce and ensuring that health workers are not linked in any way with the transmission of the HBV in the general population. Hepatitis B and C are global health problems mostly in the developing countries. Besides other modes of infection, occupational exposure of health care workers to infected blood remains one of the major modes of infection and risk factor.

### Recommendations

There is need to educate our HCWs through a well-organized infection control programme,spreading awareness and education of infection control measures, diseases transmissions, post exposure prophylaxis and on benefits of vaccines and other preventive ways so that a change in attitude can be successfully achieved. This should be then ensured by making continuous availability of vaccine by the health institution by bearing the cost for vaccinating and organizing vaccination programme to achieve 100% coverage with administrative support as multidisciplinary approach is required. Another effective way worth following is that vaccination of health care workers, medical and paramedical students can be made mandatory at the time of entry in service or studies.

As persistence of HBV infection has grave consequences and no satisfactory treatment is available for, the Nursing staff should take all the preventive measures to save themselves of this menace. All the staff should be given vaccination for HBV at the start of their career. All healthcare workers should be educated regarding universal precautions.

